# Biomedical Text Embeddings Weakly Capture Chromatin Accessibility Similarity: A Large-Scale Measurement Study

**DOI:** 10.64898/2026.11.23.768614

**Authors:** Joshua Song, Brandon Lukas, Yang Dai

## Abstract

Text metadata provides the most readily available description of public chromatin accessibility data, but it remains unclear how well it reflects similarity between the underlying chromatin profiles. We systematically compared pairwise cosine similarity matrices derived from 12 embedding types across 32,245 human assay for transposase-accessible chromatin with sequencing (ATAC-seq) experiments obtained from ChIP-Atlas. The embeddings were two chromatin representations (Atacformer and regulatory potential, RP), three biomedical language models and one general-domain model applied to full metadata and to cell-type names alone, term frequency-inverse document frequency (TF-IDF), and a random baseline. We measured agreement using Spearman rank correlation, Mantel permutation tests, and top-*k* neighborhood overlap in the similarity matrices. Across the full dataset, text-derived similarities correlated with Atacformer similarities at *ρ* = 0.20 to 0.30. This text-Atacformer agreement was largely driven by experiments with relatively few called peaks, which were enriched for promoter regions (median 69 to 75% within 1 kb of a transcription start site for experiments with 100 to 10,000 peaks), resulting in similar chromatin representations regardless of cell type. Among the 17,232 experiments with at least 5,000 peaks, the text-Atacformer correlations dropped substantially (*ρ* = 0.011 to 0.016), with less than 0.1% of top-5 neighbors shared. Atacformer itself barely tracked cell type in this subset, with only 1.3% of an experiment’s top-5 neighbors sharing its cell-type label (0.8% expected by chance). In contrast, text-RP agreement was weak but persisted (*ρ* = 0.06 to 0.10; 28 to 35% of top-5 neighbors shared), suggesting that both capture coarse cell-type identity. Most of the text-derived similarity signal in the full dataset came from sample titles, which mainly identify the submission and sequencing platform. Notably, TF-IDF matched or exceeded the language models in every correlation comparison. These findings indicate that text metadata is at best a coarse proxy for chromatin accessibility similarity, and highlight the importance of controlling for peak count when comparing chromatin representations across public repositories.

## 1 Introduction

The rapid growth of epigenomics databases, such as ChIP-Atlas [1], Encyclopedia of DNA Elements (ENCODE) [2], and Roadmap Epigenomics [3], has produced tens of thousands of ATAC-seq experiments across diverse human cell types and conditions. These growing repositories provide unprecedented opportunities for systematic exploration of chromatin regulatory landscapes, discovery of biological knowledge, and development of AI-ready epigenomic datasets. Organizing, retrieving, and comparing these experiments requires similarity measures that reflect the underlying biology. Text metadata, including cell-type annotations, cell-type descriptions, and experiment titles, offers a readily accessible representation for indexing and searching these datasets. However, it remains unclear whether similarities derived from text metadata faithfully reflect the underlying chromatin accessibility profiles.

This question has important implications. If text embeddings derived from experimental metadata closely approximate chromatin-based similarity measures, such as those derived from Atacformer [4] or regulatory potential (RP) [5], they could enable natural-language retrieval and annotation of ATAC-seq data without requiring processing of raw chromatin accessibility profiles. Such capabilities would facilitate the integration and reuse of public epigenomic data in AI-driven research. If agreement is limited, the results instead point to fundamental limits on what text can capture and motivate investment in chromatin-specific representation learning.

Recent advances in biomedical natural language processing have produced powerful text encoders pretrained on large biomedical corpora. PubMedBERT [6] is pretrained from scratch on PubMed abstracts and full-text articles. SapBERT [7] and MedCPT [8] are both initialized from a pretrained PubMedBERT checkpoint and further fine-tuned for different objectives: SapBERT through self-alignment on Unified Medical Language System (UMLS) synonym pairs, and MedCPT through contrastive learning on PubMed search logs. Although these models have demonstrated strong performance in biomedical entity recognition, relation extraction, and information retrieval, whether their representations capture the biological relationships encoded in chromatin accessibility data has not been systematically evaluated.

In contrast to text-based representations, chromatin-derived embeddings directly capture features of the underlying accessibility profiles. Atacformer [4] is a transformer trained on ATAC-seq data with a triplet margin loss so that its embeddings cluster by cell type and chromatin state. RP [5] provides a simpler, gene-centered alternative in which each gene is scored by the peaks near its transcription start site (TSS), weighted by their distance from it.

Here, we systematically evaluate pairwise cosine similarity matrices constructed from the 12 embedding types (two chromatin representations, four language models applied to full metadata and to cell-type names, term frequency-inverse document frequency (TF-IDF), and a random baseline) across 32,245 curated human ATAC-seq experiments, treating both agreement and disagreement as informative findings. TF-IDF represents each metadata string by its word frequencies alone, so it serves as a purely lexical baseline for testing whether the pretrained language models capture anything beyond shared vocabulary. Since public repositories mix experiments of very different quality, we further examine how the number of called peaks affects representation similarity and focus our primary analysis on the 17,232 experiments with at least 5,000 peaks.

The key findings are:

- **Peak count strongly shapes similarity in both chromatin representations**. Experiments with relatively few peaks consist mostly of promoter peaks, and they inflate apparent text-chromatin agreement on the full dataset.
- **Text-derived similarities show limited agreement with chromatin-based representations**. In experiments with at least 5,000 peaks, text-derived similarities showed negligible agreement with Atacformer and only weak agreement with RP, mostly through coarse cell-type identity.
- **Pretrained language models offer little improvement over TF-IDF**. TF-IDF performed comparably to all four language models. On the full dataset, most of the text-derived signal comes from sample titles that reflect data provenance more than biological characteristics.

Together, these findings highlight the limitations of text embeddings of metadata as proxies for chromatin accessibility similarity and underscore the need for biologically grounded representations to support reliable data retrieval, integration, and knowledge discovery in epigenomic repositories.

## 2 Methods

Fig. 1 summarizes the study design. Chromatin-based embeddings (Atacformer and RP) and text-based embeddings (language models, TF-IDF, and a random baseline) are computed for each experiment, converted into pairwise similarity matrices at the cell-type and experiment levels, and compared using the metrics described in Section 2.5.

**Figure 1.**
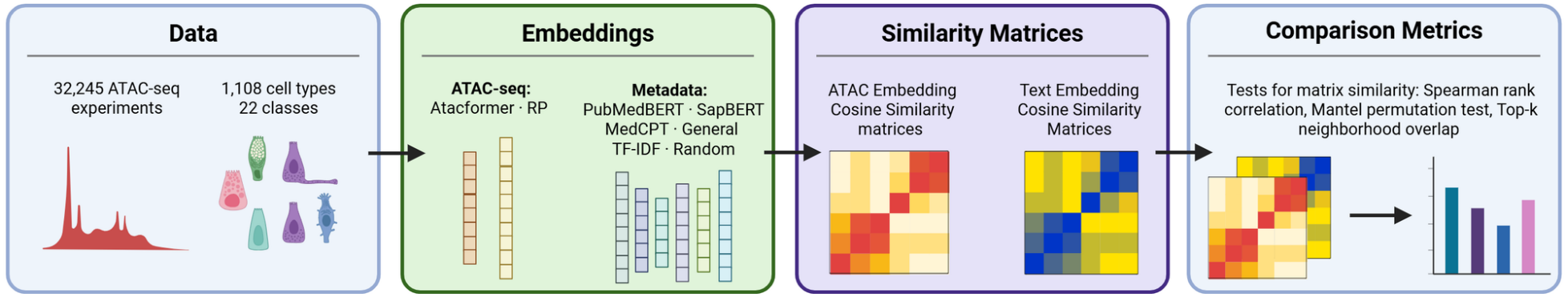
Study design. ATAC-seq experiments are embedded with chromatin-based models (Atacformer, RP), text encoders (biomedical and general-domain language models, TF-IDF), and a random baseline. Pairwise cosine similarity matrices are computed at the cell-type and experiment levels and compared using Spearman rank correlation, Mantel permutation tests, and top-*k* neighborhood overlap. Created in https://BioRender.com.

### 2.1 Dataset

We obtained 48,822 human ATAC-seq experiments from ChIP-Atlas (hg38 assembly) [1]. We removed 16,567 experiments with corrupted or empty BED files and 10 experiments producing zero regulatory potential signal near gene TSSs, leaving a curated set of 32,245 experiments spanning 1,108 cell types.

We used the ChIP-Atlas peak calls at *q <* 10^*−*10^. The number of peaks per experiment ranges from 1 to 181,134 (median 6,287; interquartile range 780 to 19,836), and 8,964 experiments have fewer than 1,000 peaks. Because experiments with few peaks retain mostly promoter peaks (Section 3.1), our primary analysis uses the 17,232 experiments with at least 5,000 peaks, which we call *well-covered* experiments. We use peak count as a proxy for sequencing depth and library quality, which were not part of the metadata we used. Results on all 32,245 experiments (the *full dataset*) are reported for comparison.

### 2.2 Atacformer Embeddings

We encoded each experiment using `databio/atacformer-base-hg38` (AtacformerForCellClustering) [4], a transformer trained with a triplet margin loss. Peak BED files were tokenized using the gtars Tokenizer with a vocabulary of 890K genomic regions. Up to 1,024 randomly sampled peak tokens were mean-pooled to produce a 192-dimensional embedding per experiment.

### 2.3 Regulatory Potential Embeddings

We computed RP scores for each experiment using a LISA2-style exponential decay model [5]. Each peak’s contribution to a gene is weighted by *w*(*d*) = *e*^*−d/*10kb^, where *d* is the distance to the gene’s transcription start site (TSS), and peaks overlapping annotated exons receive a fixed weight of 1.0 instead. Summing over all peaks gives a per-gene RP score, and stacking these scores over all hg38 autosomal gene TSSs maps each experiment to a 40,772-dimensional RP vector.

### 2.4 Text Embeddings

For each experiment, we built a metadata string by concatenating the available fields in the form

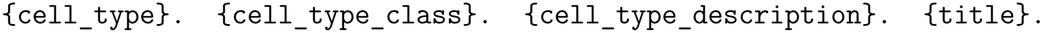

The description is the ChIP-Atlas cell-type description (for example, primary tissue and diagnosis), and the title is the experiment title from the sequencing archive record. We embedded this string with four encoders, each producing 768-dimensional embeddings. Three are biomedical models: PubMedBERT [6] (`microsoft/BiomedNLP-BiomedBERT-base-uncased-abstract-fullt ext`, mean pooling), SapBERT [7] (`cambridgeltl/SapBERT-from-PubMedBERT-fulltext`, classification token (CLS) pooling), and MedCPT [8] (`ncbi/MedCPT-Query-Encoder`, mean pooling). The fourth is a general-domain sentence encoder [9] (`sentence-transformers/all-m pnet-base-v2`, mean pooling), which we call General. We refer to the four encoders as language models (LMs). To isolate the contribution of cell-type identity, we also encoded the cell-type name alone with all four models (name-only variants). We additionally computed TF-IDF embeddings (top 2,048 features, sublinear term frequency, 2,048-dimensional) and a random Gaussian baseline (192-dimensional, seed 42).

### 2.5 Similarity Matrices and Comparison Metrics

For each embedding type, we constructed pairwise cosine similarity matrices at two levels, after subtracting a corpus mean vector from every embedding. Corpus centering corrects for anisotropy in transformer mean-pooled representations. Without it, embeddings tend to occupy a narrow cone, which inflates all cosine similarities and hides relative differences between pairs. For each embedding model, the corpus mean vector was the element-wise mean of 10,000 individual experiment embeddings, namely the first 10,000 files of that model’s embedding directory in file-system order. This is a fixed subset rather than a random sample. For every model except Atacformer, all 10,000 belong to the curated set; for Atacformer, 3 come from experiments excluded during curation. The mean was computed from individual experiments rather than from cell-type means, so the same vector is used at both levels below and does not depend on the level of aggregation.

#### Cell-type level

Experiment embeddings were mean-pooled within each cell type to give one vector per cell type, and the corpus mean vector was subtracted from each cell-type vector. For every pair of cell types (*i, j*), the cosine similarity between their centered vectors was stored in entry (*i, j*), giving a symmetric matrix with ones on the diagonal. For the full dataset, this matrix is 1,108 *×* 1,108, with one row and one column for each of the 1,108 cell types annotated in ChIP-Atlas.

#### Experiment level

The same corpus mean vector was subtracted from each individual experiment embedding, and the cosine similarity between every pair of experiments (*i, j*) was stored in entry (*i, j*) (32,245 *×* 32,245 for the full dataset). We call entry (*i, j*) of a representation’s matrix the similarity of experiments *i* and *j* under that representation, for example their Atacformer similarity or RP similarity. For any of the text embeddings, we call it their text-derived similarity.

We compared each text-derived similarity matrix with each chromatin reference (Atacformer and RP) in three ways, and refer to the result as text-Atacformer or text-RP agreement. The main measure was the Spearman rank correlation (*ρ*) between the upper-triangle entries of the two matrices, with bootstrap 95% confidence intervals (CIs) from 1,000 resamples of cell types or experiments (50,000 randomly sampled pairs per resample). At the cell-type level, we also ran one-sided Mantel permutation tests (1,000 permutations). Finally, we measured top-*k* neighborhood overlap at the experiment level. For two representations A and B, we took each experiment’s *k* most similar experiments in A’s similarity matrix and its *k* most similar experiments in B’s similarity matrix, and recorded the fraction of experiments shared between the two lists. The top-*k* overlap between A and B is this fraction averaged over all experiments. We report it for each text embedding against each chromatin reference, for Atacformer against RP, and, in Section 2.7, for each LM against TF-IDF, with *k ∈ {*5, 10, 20, 50*}*.

For the primary analysis, cell-type vectors were recomputed from well-covered experiments only (954 cell types), and experiment-level matrices were restricted to the 17,232 well-covered experiments. The corpus mean vectors used for centering were the same as above, and experiment-level correlations were computed on 2 *×* 10^6^ randomly sampled pairs.

### 2.6 Technical Covariates and Sensitivity Analyses

For each experiment, we recorded the number of peaks in its BED file and the fraction of peaks whose midpoint lies within 1 kb of an annotated RefSeq TSS (the hg38 annotation used for RP). For a pair of experiments with *n*_*i*_ and *n* _*j*_ peaks, we defined *depth match* as *−*| log_10_ *n*_*i*_ *−* log_10_ *n* _*j*_| and *mean depth* as (log_10_ *n*_*i*_ + log_10_ *n* _*j*_)*/*2.

We used these covariates in four sensitivity analyses, each testing whether an observed agreement depends on a technical factor rather than on biology. All four were run on the full dataset: (i) Spearman correlations within peak-count strata, using pairs in which both experiments have fewer than 1,000, 1,000 to 5,000, 5,000 to 10,000, or at least 10,000 peaks; (ii) partial Spearman correlations (Section 2.7) controlling for depth match and mean depth, and separately for indicators of shared cell type and shared cell-type class, computed by regressing the covariate ranks out of each variable’s ranks; (iii) a title-only TF-IDF embedding with the same settings, fit on the title field alone and corpus-centered, to isolate the contribution of sample titles; and (iv) recomputation of the Atacformer, RP, full-text LM, and TF-IDF similarities without corpus centering, to test whether the results depend on centering.

Separately from these sensitivity analyses, we quantified the statistical uncertainty of every partial correlation reported in this study, including those in Section 2.7, with an experiment-level block bootstrap (1,000 resamples of experiments, 200,000 pairs each).

### 2.7 Comparing Language Models with TF-IDF

To assess whether the language model (LM) embeddings capture information related to chromatin similarity beyond vocabulary overlap, we compared the full-text LM embeddings with TF-IDF representations in three ways.

#### Pairwise similarity scatter

On the full dataset, we sampled 25,000 random experiment pairs, plotted TF-IDF similarity against LM similarity, and colored each point by Atacformer similarity. Points above the diagonal are pairs where the LM score is higher than the TF-IDF score. We also computed the Spearman correlation between the TF-IDF and LM similarities on the same pairs.

#### Partial correlation

We computed the partial Spearman correlation of each LM similarity matrix with each chromatin reference, controlling for TF-IDF (TF). For Atacformer (AC), this is

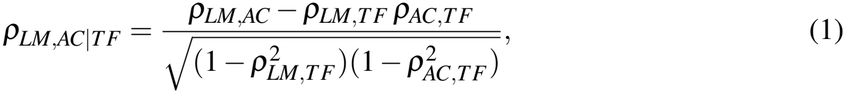

where *ρ*_*X,Y*_ is the Spearman correlation between the upper-triangle entries of matrices *X* and *Y* , estimated on 2 *×* 10^6^ random pairs (out of about 148 million pairs for well-covered experiments and 520 million for the full dataset). A positive value means that the LM similarity is associated with chromatin similarity beyond what TF-IDF explains. A value near zero or below means that TF-IDF accounts for the LM’s agreement with the reference.

#### Top-k overlap with TF-IDF

For each LM, we computed its top-*k* overlap with TF-IDF (Section 2.5), that is, the mean fraction of its top-*k* nearest neighbors that are also top-*k* TF-IDF neighbors (*k ∈ {*5, 10, 20, 50*}* on the full dataset and *k* = 5 in well-covered experiments). High overlap means that the LM’s local structure largely reproduces that of TF-IDF.

## 3 Results

### 3.1 Peak Count Inflates Apparent Text-Chromatin Agreement

Experiments with few peaks consist mostly of promoter peaks (Fig. 2a). In every peak-count bin between 100 and 10,000 peaks, the median experiment had 69 to 75% of its peaks within 1 kb of a TSS. The fraction progressively decreased with increasing peak count, to 57% in experiments with 10,000 to 20,000 peaks, 39% in those with 20,000 to 50,000 peaks, and 24% in those with more than 50,000 peaks. The 1,689 experiments with fewer than 100 peaks were an exception, exhibiting a median promoter-proximal peak fraction of 42%. Because promoters of broadly expressed genes are accessible in most cell types, experiments that retain mainly promoter peaks tend to resemble one another regardless of cell-type identity.

**Figure 2.**
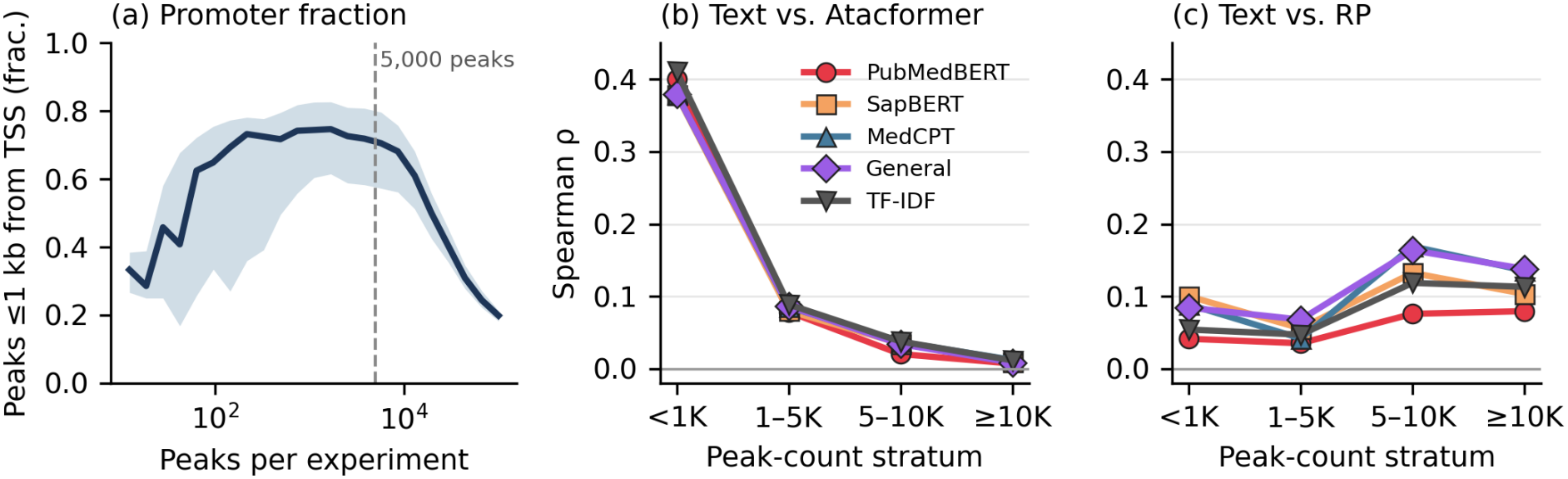
Peak count and text-chromatin agreement on the full dataset. (a) Fraction of each experiment’s peaks within 1 kb of an annotated TSS against its number of peaks (median and interquartile range in log-spaced bins containing at least 30 experiments). The dashed line marks the 5,000-peak threshold used for the primary analysis. (b, c) Spearman correlation between text-derived similarity and Atacformer (b) or RP (c) similarity, computed separately for pairs in which both experiments fall in the same peak-count stratum (2 *×* 10^6^ random pairs in total).

Peak count was also strongly associated with similarity in both chromatin representations. On the full dataset, the Atacformer similarity of two experiments (the cosine similarity of their Atacformer embeddings) correlated with their depth match (Section 2.6) at *ρ* = 0.66, and their RP similarity correlated with depth match at *ρ* = 0.59. Both associations were stronger than any correlation observed for a text embedding. After controlling for depth match and mean depth, the experiment-level correlation between Atacformer and RP decreased from 0.22 to *−*0.02. Peak count matters within cell types as well. Across 406 cell types with at least 10 experiments, mean within-cell-type Atacformer similarity was lower in cell types with a wider spread of peak counts (Spearman *ρ* = *−*0.77 against the standard deviation of log_10_ peak count; RP: *ρ* = *−*0.71). These findings highlight peak count as an important factor influencing chromatin-based similarity, even among experiments sharing the same cell-type annotation.

Text-Atacformer agreement on the full dataset was largely driven by low-peak experiments (Fig. 2b). When both experiments in a pair had fewer than 1,000 peaks, correlations between text-derived and Atacformer similarities were highest, at *ρ* = 0.38 to 0.41. The correlation dropped to 0.08 to 0.09 at 1,000 to 5,000 peaks, 0.02 to 0.04 at 5,000 to 10,000 peaks, and 0.007 to 0.012 at 10,000 peaks or more. Agreement with RP behaved differently (Fig. 2c). It was lowest in the two low-peak strata (*ρ* = 0.04 to 0.10) and higher in the two well-covered strata (*ρ* = 0.08 to 0.17). These contrasting patterns suggest that the relationship between text-derived and chromatin-derived similarities depends strongly on peak count and differs substantially between Atacformer and RP.

Within the metadata, most of the signal comes from sample titles. On the same 2 *×* 10^6^ pairs, a TF-IDF embedding of the title field alone correlated with Atacformer as strongly as PubMedBERT applied to the full metadata string (*ρ* = 0.275 for both). Within cell types, where the other fields are mostly constant, the title-only embedding exceeded every full-text model (*ρ* = 0.24 vs. 0.20 to 0.22), and controlling for title similarity reduced the within-cell-type correlations of the text embeddings with Atacformer to 0.04 to 0.06. The titles are archive sample names such as “GSM3381886: F 2 p; Homo sapiens; ATAC-seq” or “Illumina HiSeq 2500 sequencing of SAMD00123421”. Many of them name only the submission or the sequencing platform, so the shared signal most likely reflects data provenance. Provenance is also tied to peak count. Controlling for depth match and mean depth reduced the title-only correlation with Atacformer from 0.275 to 0.092. Cell-type labels, by comparison, explained little of the full-dataset correlation. Controlling for shared cell type and shared cell-type class reduced PubMedBERT’s correlation with Atacformer only from 0.275 to 0.254. These results suggest that the apparent agreement between text-derived and Atacformer similarities is more strongly associated with sample provenance and peak-count-related variation than with annotated cell-type identity.

We therefore restricted our primary analysis to well-covered experiments with at least 5,000 peaks. The threshold was selected pragmatically. Although experiments with 5,000 to 10,000 peaks remained enriched for promoters (Fig. 2a), experiment-level analyses using a more stringent threshold of 10,000 peaks yielded consistent conclusions. These results support the robustness of our findings to the choice of peak-count threshold.

### 3.2 Text Weakly Tracks RP but Not Atacformer in Well-Covered Experiments

To qualitatively assess whether the different representations capture biologically meaningful relationships, we generated cell-type-level embeddings for 34 representative cell types by averaging the embeddings across their corresponding experiments (Fig. 3; UMAP projections in Fig. S7). Atacformer and RP showed coherent lineage structure, with immune, neural, and cancer cell types forming distinct groups, whereas the text embeddings showed less clearly resolved biological organization.

**Figure 3.**
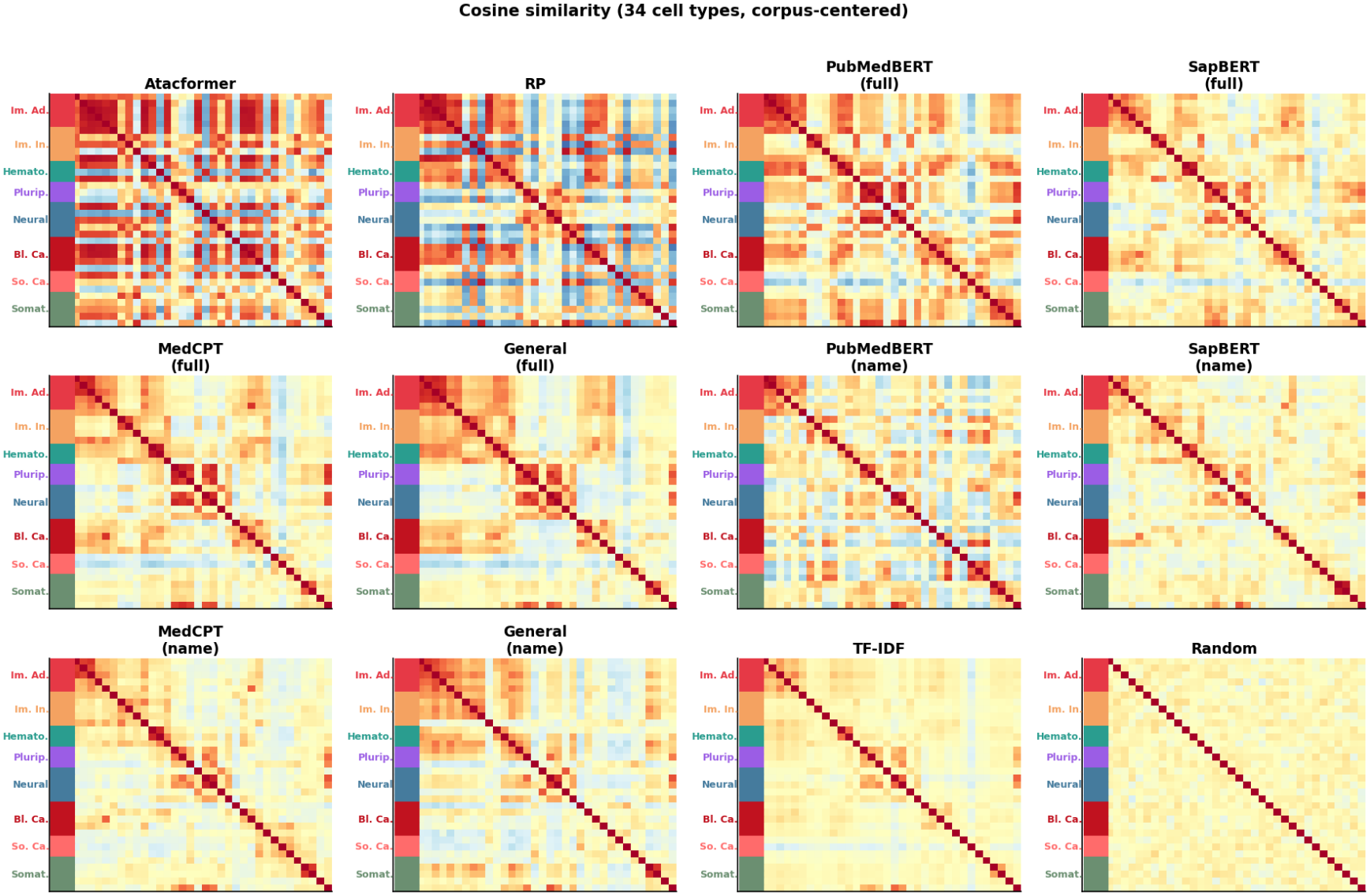
Cosine similarity between cell-type mean embeddings for 34 representative cell types chosen by hand to span eight lineage groups, shown for all 12 embedding conditions. Each cell-type embedding is the mean over all of its experiments. Rows and columns are ordered by lineage, marked by the colored strip: Im. Ad., immune adaptive; Im. In., immune innate; Hemato., hematopoietic; Plurip., pluripotent; Bl. Ca., blood cancer; So. Ca., solid cancer; Somat., somatic. Colors run from *−* 1 (blue) to 1 (red). For this figure, the corpus mean vector was computed from the first 5,000 rather than 10,000 embedding files (Section 2.5). Uniform Manifold Approximation and Projection (UMAP) projections of the same embeddings are shown in Fig. S7.

Table 1 summarizes agreement among embedding representations across the 17,232 well-covered experiments spanning 954 cell types. Text embeddings showed negligible agreement with Atacformer. At the cell-type level, the correlations of the full-text LMs and TF-IDF with Atacformer were zero or slightly negative (*ρ* = *−*0.074 to *−*0.023; one-sided Mantel *p >* 0.98). At the experiment level, they were close to zero (*ρ* = 0.011 to 0.016), compared with 0.20 to 0.30 on the full dataset. Agreement with RP was weak but consistent. The full-text LMs and TF-IDF correlated with RP at *ρ* = 0.038 to 0.068 at the cell-type level (Mantel *p* = 0.001) and at 0.061 to 0.098 at the experiment level (full-dataset values in Table 1 and Fig. S1). Atacformer and RP agreed with each other at the cell-type level (*ρ* = 0.365) but only modestly at the experiment level (*ρ* = 0.107).

**Table 1.** Spearman correlation (*ρ*) between each embedding’s similarity matrix and the Atacformer and RP similarity matrices. Primary analysis: well-covered experiments (*≥* 5,000 peaks; 954 cell types and 17,232 experiments), with 95% bootstrap CIs at the experiment level (1,000 resamples of experiments, 50,000 pairs each). Full: all 32,245 experiments. Significance markers (one-sided Mantel test for positive association, 1,000 permutations) are given for the cell-type level only.

|  | vs. Atacformer |  |  | vs. RP |  |  |
|---|---|---|---|---|---|---|
|  | ≥5,000 peaks |  | Full | ≥5,000 peaks |  | Full |
| Embedding | Cell type | Experiment [95% CI] | Experiment | Cell type | Experiment [95% CI] | Experiment |
| Atacformer / RP | 0.365<sup>***</sup> | 0.107 [0.097, 0.116] | 0.216 | 0.365<sup>***</sup> | 0.107 [0.097, 0.117] | 0.216 |
| PubMedBERT (full text) | −0.030 | 0.011 [0.001, 0.020] | 0.276 | 0.038<sup>***</sup> | 0.061 [0.051, 0.072] | 0.092 |
| SapBERT (full text) | −0.023 | 0.013 [0.004, 0.022] | 0.248 | 0.066<sup>***</sup> | 0.078 [0.069, 0.088] | 0.074 |
| MedCPT (full text) | −0.074 | 0.016 [0.006, 0.025] | 0.229 | 0.062<sup>***</sup> | 0.098 [0.087, 0.106] | 0.090 |
| General (full text) | −0.053 | 0.014 [0.005, 0.023] | 0.196 | 0.068<sup>***</sup> | 0.095 [0.085, 0.104] | 0.088 |
| TF-IDF | −0.057 | 0.016 [0.007, 0.025] | 0.300 | 0.065<sup>***</sup> | 0.091 [0.081, 0.101] | 0.089 |
| PubMedBERT (name only) | −0.071 | 0.005 [−0.004, 0.013] | 0.090 | 0.003 | 0.035 [0.026, 0.044] | 0.018 |
| SapBERT (name only) | −0.101 | 0.008 [−0.001, 0.017] | 0.137 | 0.013 | 0.065 [0.056, 0.074] | 0.026 |
| MedCPT (name only) | −0.098 | 0.012 [0.002, 0.020] | 0.068 | 0.021<sup>*</sup> | 0.080 [0.070, 0.089] | 0.022 |
| General (name only) | −0.060 | 0.006 [−0.003, 0.016] | 0.090 | 0.015<sup>*</sup> | 0.039 [0.030, 0.048] | 0.010 |
| Random Gaussian | 0.003<sup>*</sup> | 0.000 [−0.008, 0.009] | 0.000 | 0.003<sup>*</sup> | 0.000 [−0.009, 0.009] | 0.000 |

<sup>\*\*\*</sup>*p* ≤ 0.001; <sup>\*</sup>*p* < 0.05. The small random-baseline correlation at the cell-type level likely reflects centering, because subtracting the corpus mean gives the near-zero means of large cell types a common direction.

In contrast, at the cell-type level, the name-only variants agreed with RP much less than the full-text models (*ρ* = 0.003 to 0.021 vs. 0.038 to 0.068), suggesting that the cell-type description and title fields provide additional information beyond the cell-type name. At the experiment level, the gap was smaller (*ρ* = 0.035 to 0.080 for the name-only variants).

### 3.3 Language Models Add Little Beyond Vocabulary Overlap

In well-covered experiments, TF-IDF performed comparably to the four LMs on every correlation in Table 1. After controlling for TF-IDF, partial correlations with Atacformer were near zero for all four LMs (*ρ*_partial_ = 0.000 to 0.006; all bootstrap 95% CIs within *−*0.004 to 0.011). Partial correlations with RP were small for MedCPT, General, and SapBERT (*ρ*_partial_ = 0.046, 0.046, and 0.037) and near zero for PubMedBERT (0.003). LM and TF-IDF similarities were strongly correlated (*ρ* = 0.52 to 0.73), and 44 to 45% of each LM’s top-5 neighbors were also TF-IDF neighbors. These results suggest that pretrained language models capture limited additional chromatin-relevant similarity beyond that represented by TF-IDF.

On the full dataset (Fig. 4; Figs. S2 to S4), PubMedBERT (*ρ*_partial_ = 0.051; 95% CI 0.048 to 0.059) and SapBERT (0.045; CI 0.039 to 0.049) kept positive partial correlations with Atacformer, whereas MedCPT (*−*0.030) and General (*−*0.042) did not. This difference holds up under experiment-level resampling, but it disappears in well-covered experiments and depends on corpus centering. Without centering, PubMedBERT, SapBERT, and MedCPT all have clearly positive partial correlations (0.14 to 0.26), and PubMedBERT (*ρ* = 0.292) has a higher raw correlation with Atacformer than TF-IDF does (*ρ* = 0.153). Therefore, we do not interpret the full-dataset differences among LMs as evidence about their pretraining or fine-tuning objectives.

**Figure 4.**
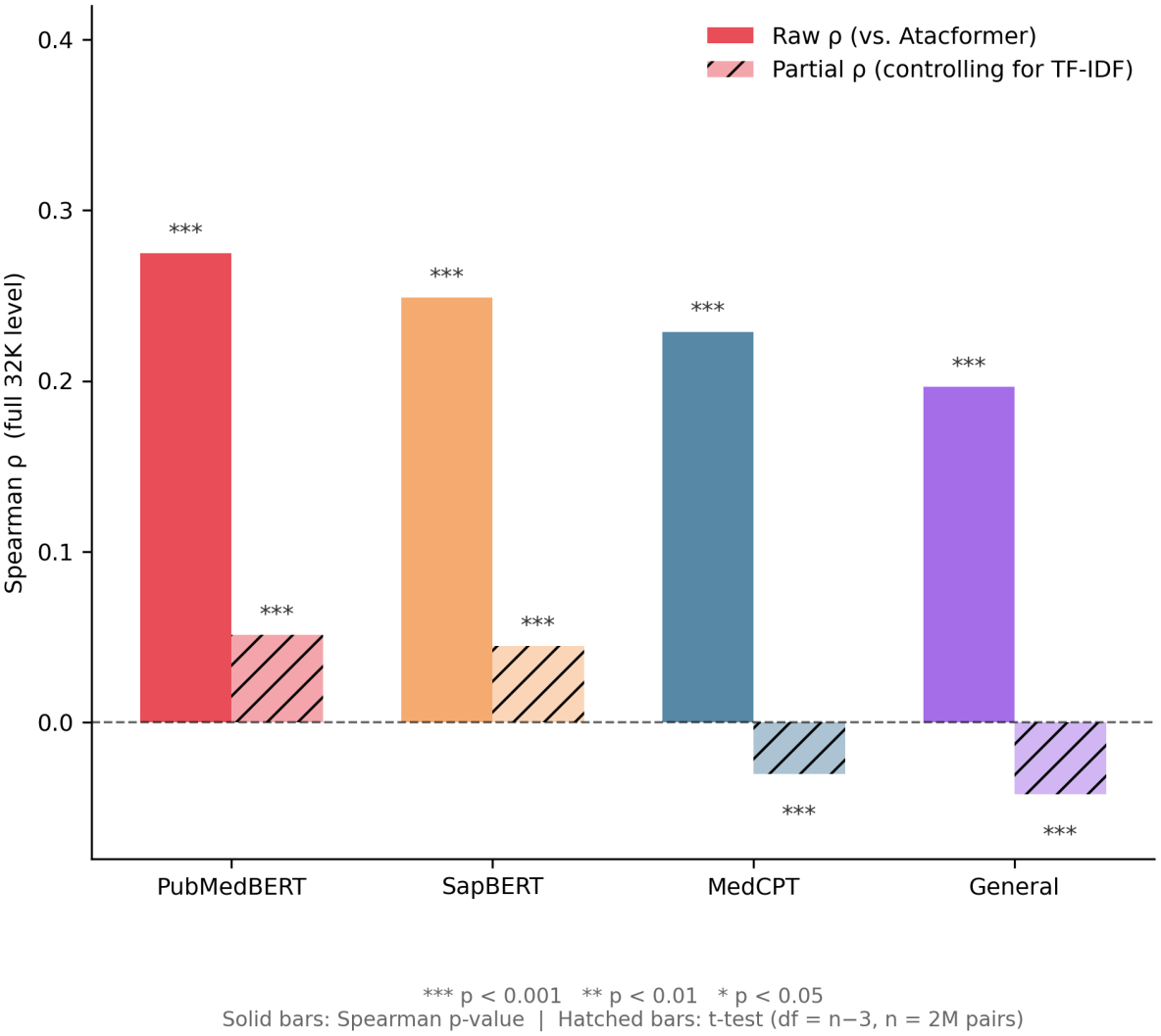
Spearman correlation with Atacformer (solid bars) and partial correlation after controlling for TF-IDF (hatched bars) for each LM on the full dataset. The significance markers in the panel come from tests that treat the sampled pairs as independent; experiment-level bootstrap CIs are given in Section 3.3. In well-covered experiments, all four partial correlations are at most 0.006.

### 3.4 Text Shares Local Structure with RP but Not Atacformer

Nearest-neighbor analyses further revealed the same differences between the text-based embeddings and the two chromatin-based representations. In well-covered experiments, the full-text LMs shared 31 to 35% of their top-5 neighbors with RP (PubMedBERT 34.7%, SapBERT 34.2%, General 32.8%, MedCPT 30.9%), TF-IDF shared 28.2%, and the name-only variants about 12%. In contrast, all text-based embeddings shared fewer than 0.1% of their top-5 neighbors with Atacformer, only slightly above the 0.03% expected by chance. Notably, Atacformer and RP themselves shared only 0.1% of their nearest neighbors, indicating that the two chromatin representations capture substantially different relationships among experiments. Across the full dataset, the corresponding values were 20 to 23% for RP and 2 to 3% for Atacformer (Figs. 5 and 6).

**Figure 5.**
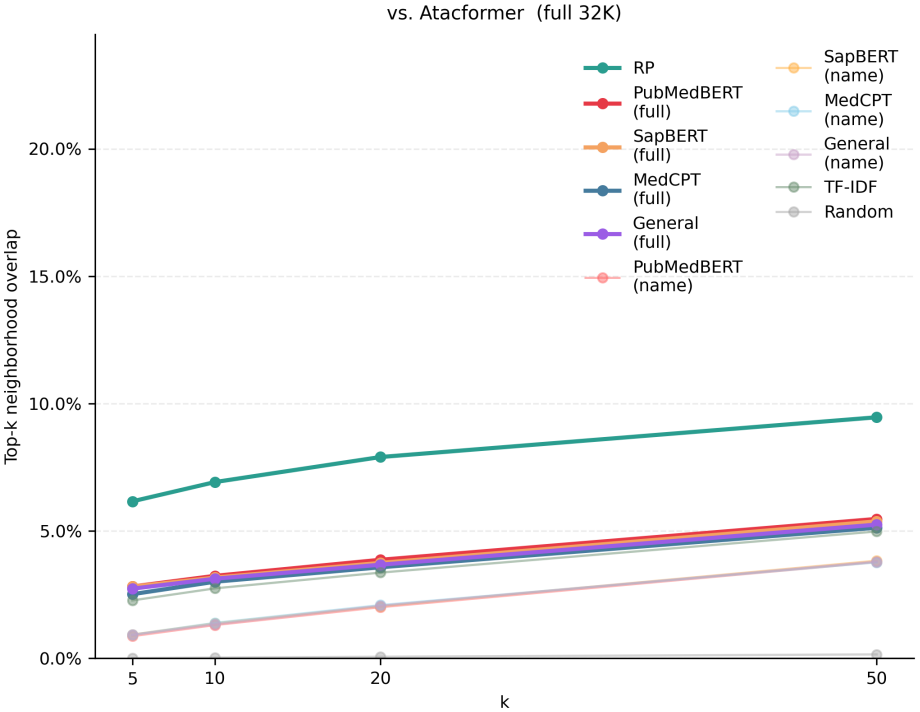
Top-*k* neighborhood overlap with Atacformer on the full dataset. Full-text models share about 2 to 3% of nearest neighbors with Atacformer at *k* = 5 (less than 0.1% in well-covered experiments).

**Figure 6.**
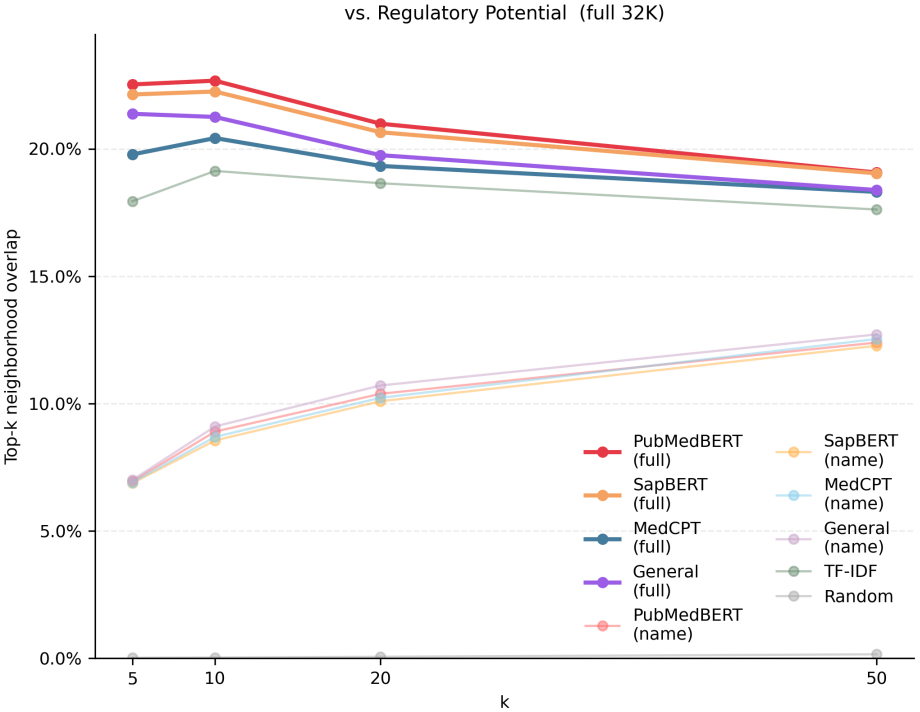
Top-*k* neighborhood overlap with RP on the full dataset. Full-text LMs share about 20 to 23% of nearest neighbors with RP at *k* = 5 (31 to 35% in well-covered experiments).

Differences in cell-type neighborhood purity provided further insight into this divergence. In well-covered experiments, 76% of an experiment’s top-5 RP neighbors shared its cell-type label (93 to 96% for the full-text embeddings), compared with only 1.3% for Atacformer, barely above the 0.8% expected by chance. On the full dataset, the values were 72% for RP and 21% for Atacformer. Within well-covered experiments, Atacformer similarity was only weakly related to depth match (*ρ* = 0.06), whereas RP similarity remained strongly related to it (*ρ* = 0.57).

We also asked how similar experiments with the same cell-type label are to one another under each representation. For each of the 796 cell types with at least two well-covered experiments, we computed the within-cell-type similarity, the mean cosine similarity over all pairs of that cell type’s experiments. Across these cell types, the median within-cell-type similarity was 0.51 for RP but only 0.17 for Atacformer, and Atacformer gave the lower value for 96.7% of cell types (paired Wilcoxon signed-rank test, *p* = 1.1 *×* 10^*−*125^). In other words, RP places experiments of the same cell type close together, whereas Atacformer does not. The same comparison on the full dataset is shown in Figs. S5 and S6.

RP thus reflects cell-type identity, which text metadata also encodes, although RP remains sensitive to peak count. In contrast, individual Atacformer embeddings of well-covered experiments carry little cell-type information. Their nearest neighbors rarely share a cell type, even though cell-type averages still agree with RP. Therefore, the low agreement between text and Atacformer is not evidence that Atacformer captures finer biological variation that text misses. One possible explanation, which we did not test, is that each Atacformer embedding averages at most 1,024 randomly sampled peaks (Section 2.2), a small and variable fraction of the peaks in a well-covered experiment.

Tables S1 and S2 list full-dataset pairs with high chromatin similarity but low TF-IDF similarity. Many of these pairs involve low-peak experiments. Every experiment in Table S2 has at most 1,438 peaks (the corpus median is 6,287), and the first pair in Table S1 (NCI-H1299 and CD4+ T cells) has 1,181 and 1,272 peaks. The remaining Atacformer pairs vary more in peak count.

## 4 Discussion

In this study, we systematically evaluated how well text-based representations of experimental metadata capture chromatin accessibility similarity across 32,245 human ATAC-seq experiments from ChIP-Atlas. We compared embeddings derived from biomedical and general-domain language models and TF-IDF against two chromatin-based representations, Atacformer and RP. Our results reveal substantial limitations in using experimental metadata as a proxy for chromatin accessibility similarity, with the observed agreement strongly influenced by peak count and sample provenance.

On the full ChIP-Atlas set, text embeddings correlated with Atacformer similarity at *ρ* = 0.20 to 0.30. However, most of this agreement came from low-peak experiments, whose promoter-dominated peak sets make them look alike in both chromatin representations, and from sample titles, which reflect data provenance and are tied to peak count. Among well-covered experiments, text embeddings barely tracked Atacformer similarity.

In contrast, text-RP agreement was weak but consistent (*ρ* = 0.06 to 0.10 at the experiment level), with 28 to 35% of top-5 neighbors shared. RP neighborhoods are strongly enriched for experiments sharing cell-type annotations, and text metadata recovers this coarse structure. These findings indicate that metadata may be useful for coarse, cell-type-level retrieval, but it should not substitute for chromatin similarity in finer-grained applications.

Language models provided limited additional information beyond simple lexical representation. In well-covered experiments, TF-IDF performed comparably to all four LMs, and partial correlations after TF-IDF adjustment were near zero for Atacformer and small for RP. Although PubMedBERT and SapBERT appeared to outperform MedCPT and General on the full dataset, these differences disappeared after restricting the analysis to well-covered experiments and were sensitive to corpus centering.

The peak-count effect matters beyond text. Similarities in both chromatin-based representations were more strongly associated with peak count than with any other variable we examined, and controlling for peak count substantially reduced the experiment-level agreement between Atacformer and RP on the full dataset. These findings identify peak-count heterogeneity as an important consideration when evaluating chromatin representations across large public repositories. Stratification or statistical adjustment for peak count, together with additional assay-quality measures, may help distinguish biologically meaningful similarities from variation associated with experimental coverage and data processing.

This study has several limitations. Atacformer and RP were used as proxies for chromatin biology without validation against an external ground truth. Both were derived from the same ChIP-Atlas peak calls and were sensitive to peak count, while the Atacformer embeddings used here carry little cell-type information for individual well-covered experiments. Our conclusions are therefore bounded by how faithfully these representations capture the biology they stand for. The metadata we used did not include sequencing depth or standard quality metrics such as the fraction of reads in peaks or TSS enrichment, so we relied on peak count as a quality proxy, with the 5,000-peak threshold selected pragmatically. We also limited the text to metadata fields available in ChIP-Atlas and did not evaluate multimodal embeddings that combine text and chromatin signals. Future work could re-embed experiments using all of their peaks, use uniformly reprocessed and quality-controlled data, and investigate whether text encoders trained to predict chromatin similarity can narrow the gap.

## Data and code availability

ATAC-seq peak calls and metadata are publicly available from ChIP-Atlas [1]. All models are public Hugging Face checkpoints, listed by identifier in Sections 2.2 and 2.4. Code for the peak-count, well-covered, and sensitivity analyses (Sections 2.6 and 3.1) and for generating the figures is available at https://github.com/jjsong23/Biological-Embedding-Project.

## Competing interests

The authors declare no competing interests.

## Acknowledgements

This work was supported by NVIDIA Corporation through its academic grant program via the provision of an NVIDIA DGX Spark system. Claude (Anthropic) [10] was used to help write analysis code and edit the manuscript. The authors reviewed all code, results, and text and take full responsibility for the content.

## Supplementary Figures and Tables

**Figure S1.**
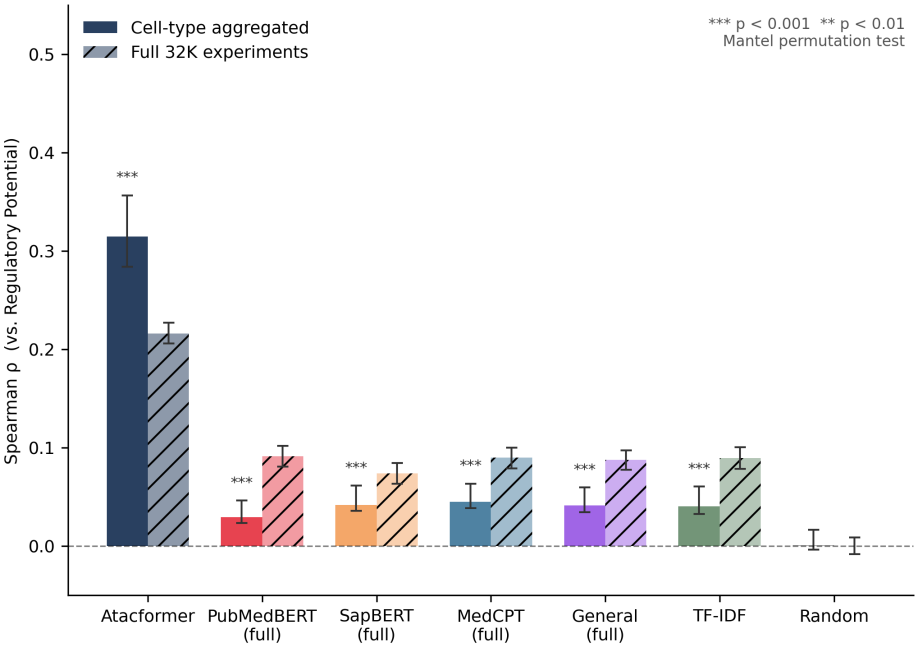
Spearman correlation between each embedding and RP on the full dataset, at the cell-type level (solid bars) and the experiment level (hatched bars); name-only variants are omitted. Error bars show 95% bootstrap CIs, and significance markers (Mantel test) apply to the cell-type bars only. Well-covered results are given in Table 1.

**Figure S2.**
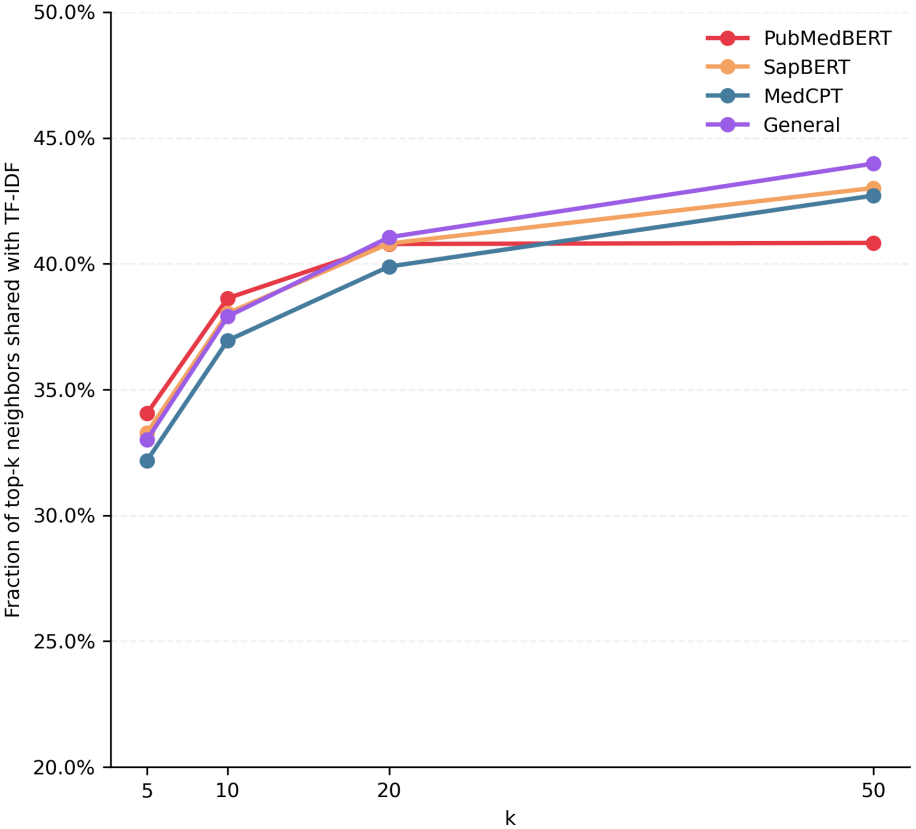
Fraction of each LM’s top-*k* nearest neighbors that are also TF-IDF neighbors on the full dataset (about 33% at *k* = 5; 44 to 45% in well-covered experiments). The *y*-axis starts at 20%.

**Figure S3.**
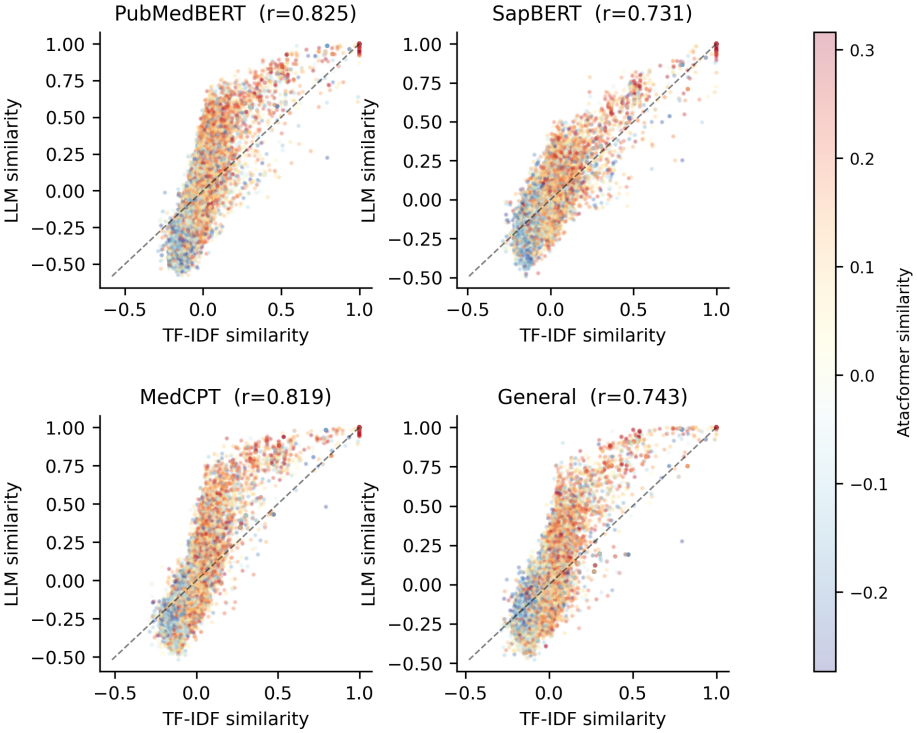
TF-IDF similarity (*x*-axis) against LM similarity (*y*-axis, labeled LLM) for 25,000 random experiment pairs from the full dataset, colored by Atacformer similarity (warmer colors indicate higher similarity). Points above the diagonal are pairs that the LM scores higher than TF-IDF. Panel titles give the Spearman correlation between the two similarities.

**Figure S4.**
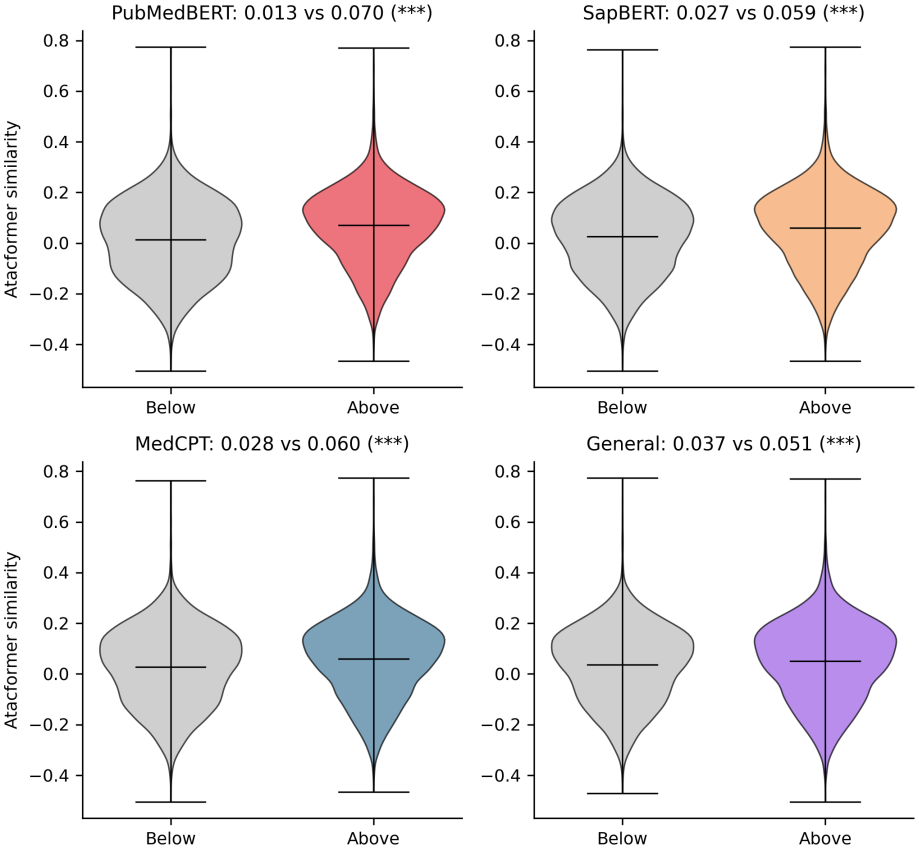
Atacformer similarity for pairs below and above the diagonal in Fig. S3 (full dataset; Welch’s *t*-test). For all four LMs, pairs above the diagonal have higher mean Atacformer similarity (PubMed-BERT: 0.070 vs. 0.013; SapBERT: 0.059 vs. 0.027; MedCPT: 0.060 vs. 0.028; General: 0.051 vs. 0.037). The corresponding effect sizes (Cohen’s *d*) are 0.38, 0.22, 0.21, and 0.09.

**Figure S5.**
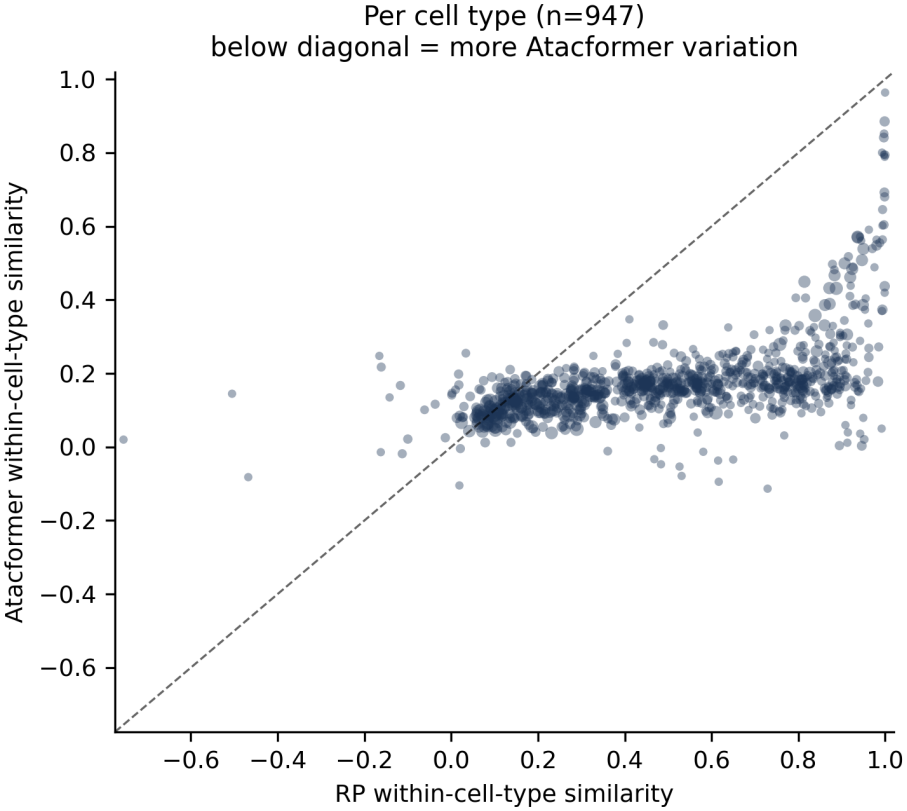
Within-cell-type similarity (mean pair-wise cosine similarity among experiments with the same cell-type label) for Atacformer against RP on the full dataset (947 cell types with at least two experiments). Points below the diagonal are cell types with lower within-cell-type similarity under Atacformer than under RP.

**Figure S6.**
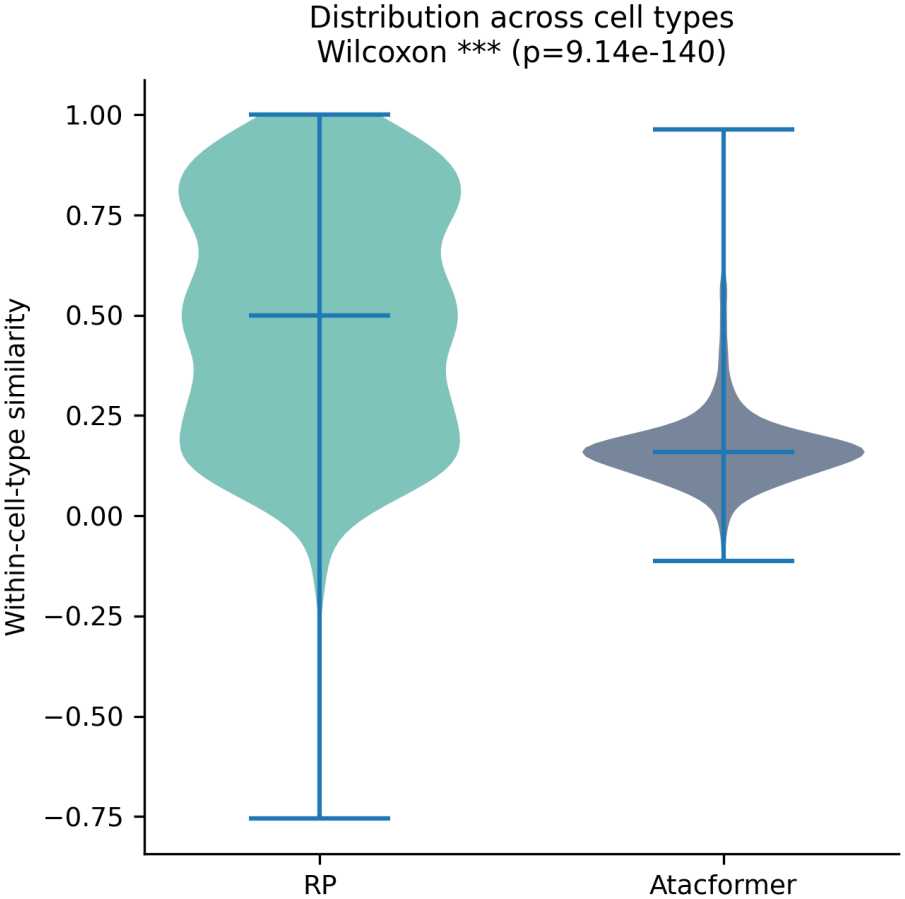
Distribution of within-cell-type similarity across cell types on the full dataset (RP median 0.50, Atacformer median 0.16; Wilcoxon signed-rank test). Within-cell-type similarity tracks the spread of peak counts within each cell type (Section 3.1).

**Figure S7.**
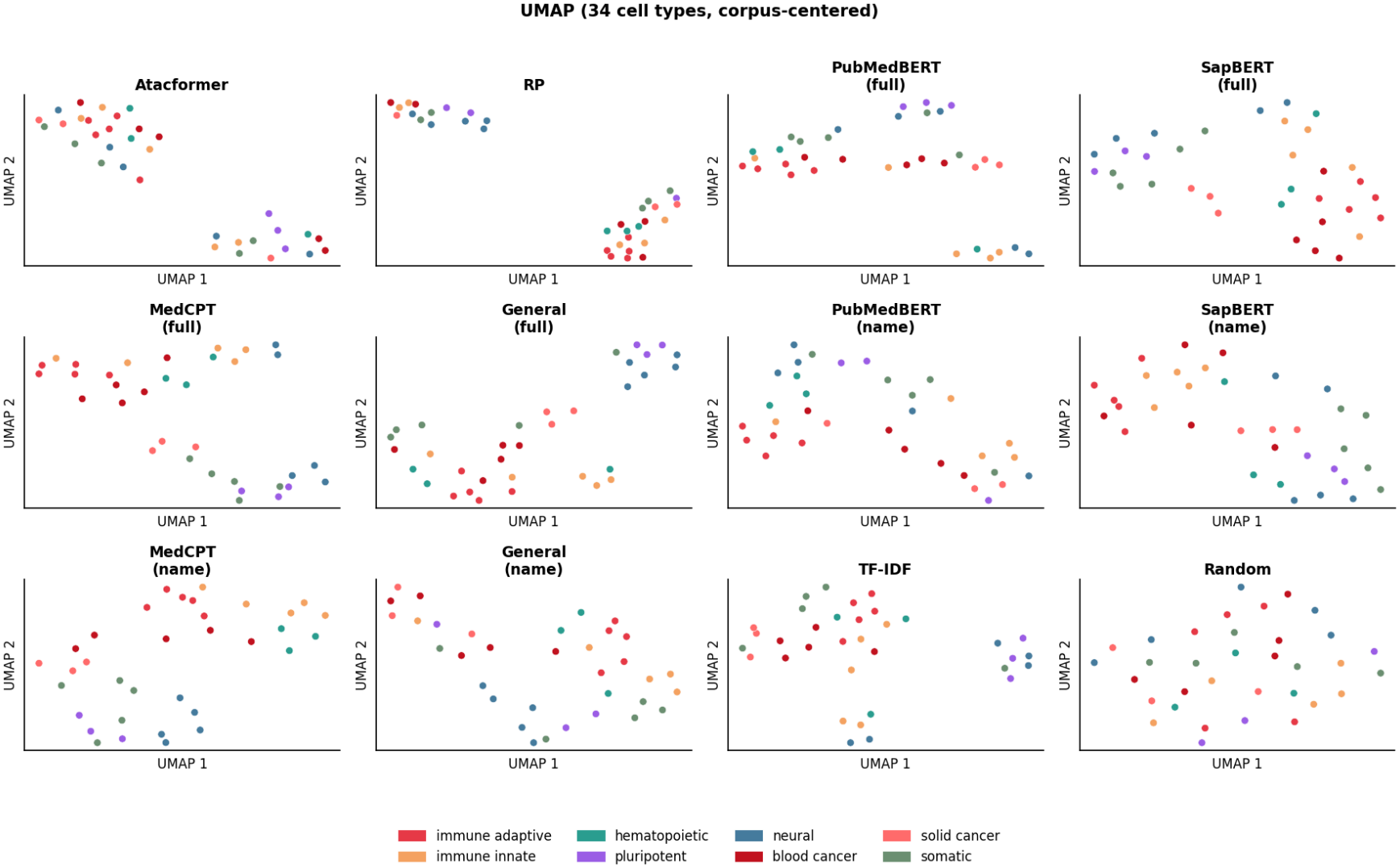
Uniform Manifold Approximation and Projection (UMAP) projections of the cell-type mean embeddings shown in Fig. 3 (34 representative cell types, all 12 embedding conditions), colored by lineage group and centered as in Fig. 3.

**Table S1.** Full-dataset experiment pairs with high Atacformer similarity (top 5%) and low TF-IDF similarity (below the median). The first pair consists of two low-peak experiments (1,181 and 1,272 peaks). Cell-type labels are given as annotated in ChIP-Atlas; “(unlabeled)” marks experiments without a cell-type annotation. iPSC, induced pluripotent stem cell.

| Cell type A | Cell type B | Atac. | TF-IDF |
|---|---|---|---|
| NCI-H1299 | CD4+ T cells | 0.374 | −0.196 |
| iPSC derived cells | PC-9 | 0.361 | −0.176 |
| NCI-H1648 | iMeLCs | 0.312 | −0.214 |
| (unlabeled) | CD4+ T cells | 0.398 | −0.147 |
| NCI-H1648 | iPSC neural cells | 0.307 | −0.204 |
| (unlabeled) | IMR-90 | 0.309 | −0.188 |
| (unlabeled) | MV-4-11 | 0.331 | −0.156 |
| (unlabeled) | Neural crest cells | 0.310 | −0.179 |
| iPSC derived cells | Jurkat | 0.318 | −0.164 |
| (unlabeled) | DND-41 | 0.316 | −0.161 |

**Table S2.** Full-dataset experiment pairs with high RP similarity and low TF-IDF similarity, selected as in Table S1. Every experiment in this table has at most 1,438 peaks, and their median promoter fraction is 72% (corpus median 57%). PBMC, peripheral blood mononuclear cells; ES, embryonic stem.

| Cell type A | Cell type B | RP | TF-IDF |
|---|---|---|---|
| iPSC derived cells | K-562 | 0.885 | −0.221 |
| NCI-H2347 | PBMC | 0.884 | −0.181 |
| RERF-LC-Ad1 | XG-6 | 0.851 | −0.224 |
| (unlabeled) | PKC19 | 0.901 | −0.171 |
| iPSC derived cells | T-47D | 0.874 | −0.185 |
| (unlabeled) | B-cell leukemia | 0.891 | −0.170 |
| NCI-H1648 | B cells | 0.859 | −0.181 |
| NCI-H1703 | CD4+ T cells | 0.851 | −0.186 |
| NCI-H1299 | CD4+ T cells | 0.833 | −0.196 |
| (unlabeled) | ES cells | 0.916 | −0.154 |

## Notes

https://chip-atlas.org/

https://chip-atlas.dbcls.jp/data/metadata/experimentList.tab

https://pypi.org/project/lisa2/

https://huggingface.co/databio/atacformer-base-hg38

https://huggingface.co/microsoft/BiomedNLP-BiomedBERT-base-uncased-abstract-fulltext

https://huggingface.co/cambridgeltl/SapBERT-from-PubMedBERT-fulltext

https://huggingface.co/ncbi/MedCPT-Query-Encoder

https://huggingface.co/sentence-transformers/all-mpnet-base-v2

https://github.com/jjsong23/Biological-Embedding-Project

### Competing Interest Statement

The authors have declared no competing interest.

## References

[1] Zhaonan Zou, Tazro Ohta, Fumihito Miura, and Shinya Oki. ChIP-Atlas 2021 update: a data-mining suite for exploring epigenomic landscapes by fully integrating ChIP-seq, ATAC-seq and Bisulfite-seq data. Nucleic Acids Research, 50(W1):W175–W182, 2022. doi: 10.1093/nar/gkac199.

[2] ENCODE Project Consortium. An integrated encyclopedia of DNA elements in the human genome. Nature, 489:57–74, 2012. doi: 10.1038/nature11247.

[3] Roadmap Epigenomics Consortium. Integrative analysis of 111 reference human epigenomes. Nature, 518:317–330, 2015. doi: 10.1038/nature14248.

[4] Nathan J. LeRoy, Guangtao Zheng, Oleksandr Khoroshevskyi, Donald R. Campbell, Aidong Zhang, and Nathan C. Sheffield. Atacformer: A transformer-based foundation model for analysis and interpretation of ATAC-seq data. bioRxiv, 2025. doi: 10.1101/2025.11.03.685753.

[5] Qian Qin, Jingyu Fan, Rongbin Zheng, Changxin Wan, Shenglin Mei, Qiu Wu, Hanfei Sun, Myles Brown, Jing Zhang, Clifford A. Meyer, and X. Shirley Liu. Lisa: inferring transcriptional regulators through integrative modeling of public chromatin accessibility and ChIP-seq data. Genome Biology, 21:32, 2020. doi: 10.1186/s13059-020-1934-6.

[6] Yu Gu, Robert Tinn, Hao Cheng, Michael Lucas, Naoto Usuyama, Xiaodong Liu, Tristan Naumann, Jianfeng Gao, and Hoifung Poon. Domain-specific language model pretraining for biomedical natural language processing. ACM Transactions on Computing for Healthcare, 3 (1):1–23, 2021. doi: 10.1145/3458754.

[7] Fangyu Liu, Ehsan Shareghi, Zaiqiao Meng, Marco Basaldella, and Nigel Collier. Self-alignment pretraining for biomedical entity representations. In Proc. NAACL, pages 4228–4238, 2021.

[8] Qiao Jin, Won Kim, Qingyu Chen, Donald C. Comeau, Lana Yeganova, W. John Wilbur, and Zhiyong Lu. MedCPT: Contrastive Pre-trained Transformers with large-scale PubMed search logs for zero-shot biomedical information retrieval. Bioinformatics, 39(11):btad651, 2023. doi: 10.1093/bioinformatics/btad651.

[9] Kaitao Song, Xu Tan, Tao Qin, Jianfeng Lu, and Tie-Yan Liu. MPNet: Masked and permuted pre-training for language understanding. In Advances in Neural Information Processing Systems, volume 33, pages 16857–16867, 2020.

[10] Anthropic. Claude. Large language model, 2026.

